# Assessing eRNAs associated with a cytokine-sensing mammary super-enhancer

**DOI:** 10.1101/174953

**Authors:** Chaochen Wang, Tyler Kuhns, Christian Reinbold, Michaela Willi, Lothar Hennighausen

## Abstract

Enhancers are transcription factor platforms that also bind RNA polymerase II and generate enhancer RNAs (eRNA). Although eRNAs have been suggested as predictors of enhancer activities and possibly key components facilitating transcription, there is little genetic evidence to support this. We address the biology of eRNAs i*n vivo* and investigate eRNA patterns, expression levels and possible functions within a mammary-specific super-enhancer that is composed of three units with distinct transcriptional capacities. We show that eRNA levels do not correspond with the activities of their respective enhancer units. However, changes in eRNA expression upon deletion of individual enhancer units reflect the change in overall super-enhancer activity. These data provide genetic evidence that eRNA levels are not a reliable readout of individual enhancers, but they predict super-enhancer activity in the absence of constituent elements.

## Introduction

Enhancers are DNA elements that augment the expression of target genes, frequently in a lineage-specific manner. In addition to transcription factors they are occupied by RNA Polymerase II (Pol II) and are typically transcribed into enhancer RNAs (eRNAs) (1-3). Several reports have shown that enhancer activation and the transcription of nearby genes directly correlates with eRNA levels (2-6), and knockdown of eRNAs in several mouse and human cell lines has been shown to result in decreased expression of specific genes (6-8).

While these experiments suggest that eRNAs play a role in activating the expression of target genes, several studies have demonstrated that the underlying DNA sequence of the enhancer, but not the associated transcript, is responsible for regulating gene expression. (9-11). Despite the controversy regarding whether eRNAs are a causal agent of enhancer-mediated activation or simply byproducts of Pol II occupation, eRNA levels are widely accepted as predictors of enhancer activity. In fact, eRNAs have been proposed to be the most accurate predictor of enhancer activity, and eRNA transcription measured by GRO-seq has been used for *de novo* enhancer prediction (12,13).

However, there is little genetic evidence to support the notion that eRNA levels predict enhancer activity. Furthermore, there are few studies investigating the relationship between super-enhancers, clusters of enhancers highly enriched for co-activators and transcription factors, and their associated eRNAs. We have addressed this question by assessing the levels of eRNAs at each of the four STAT5-occupied regions in the *Wap* super-enhancer (14). By combining RNA-seq data with previously published expression data from knockout mice harboring deletions of the constituent enhancers, we show that eRNA levels are not correlated with the ability of an enhancer to induce expression of a target gene. We also demonstrate that one of the constituent elements within the super-enhancer can serve as the transcription start site for the downstream *Ramp3* gene in the absence of the endogenous promoter.

## Results

### *Activities of constituent enhancers within the* Wap *super-enhancer do not parallel their respective eRNA levels*

The *Wap* gene, which encodes a major milk protein, is induced nearly 1,000-fold over the course of pregnancy by a STAT5-driven super-enhancer that is composed of three constituent enhancers (E1-3) and a fourth STAT5-occupied region (S4) (14) (Fig. 1A). However, the extent to which this super-enhancer is transcribed is not known. To investigate the possibility of eRNA synthesis at this locus, we first analyzed ChIP-seq data for evidence of RNA Polymerase II (Pol II) binding (Fig. 1B, C). We found that Pol II was recruited to each constituent element of the super-enhancer, suggesting that they are transcribed. The most abundant Pol II occupancy was observed at S4, which unlike E1-3 had a single Pol II peak underlying STAT5, as opposed to two peaks flanking the STAT5-occupied region. The presence of RNA-seq reads near each individual enhancer further supported the notion that this super-enhancer is transcribed.

**Figure 1.**
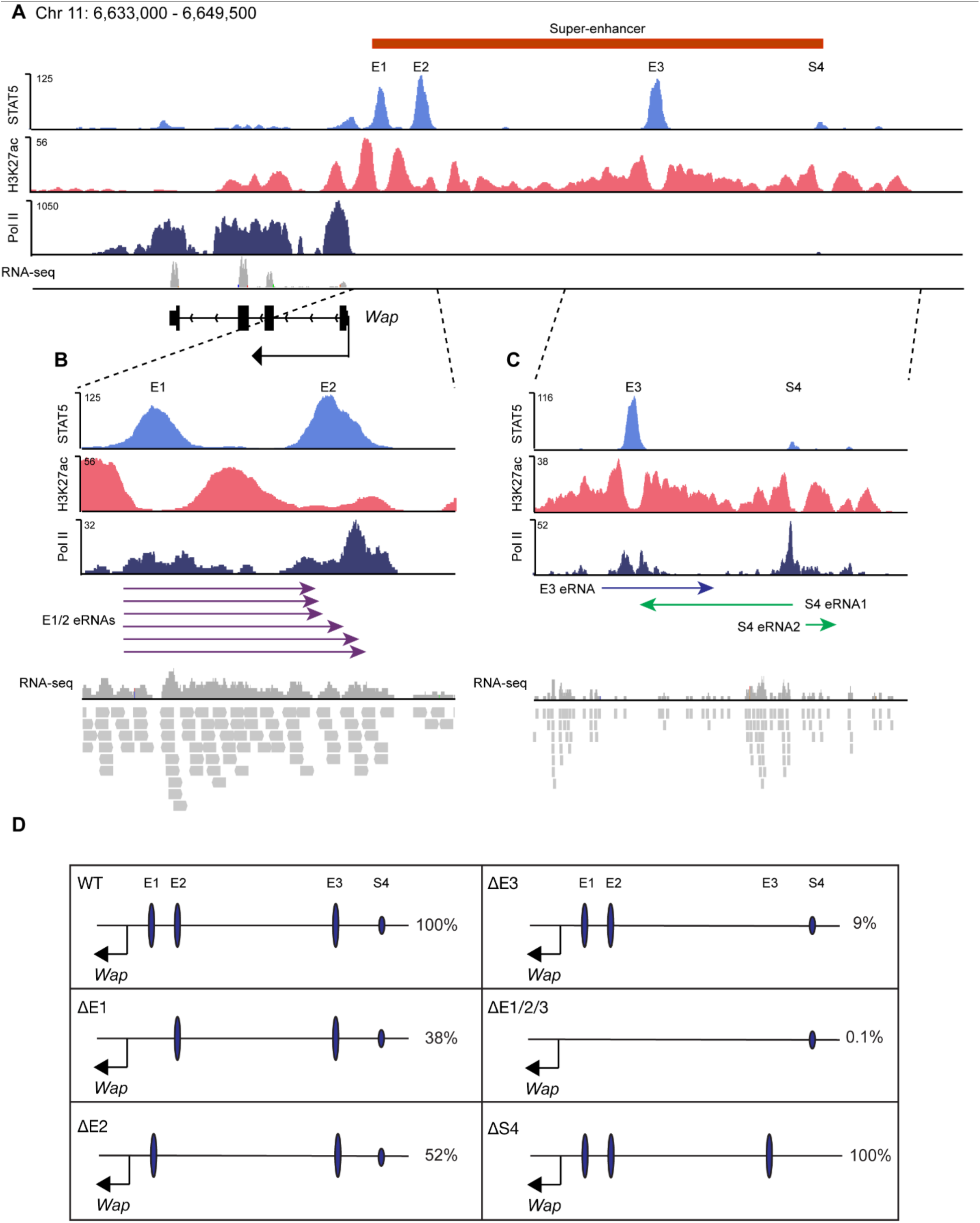
Structure of the *Wap* super-enhancer in mammary tissue at lactation day 1 (L1), the associated eRNAs, and the contribution of each constituent element to *Wap* expression at L1. (A) ChIP-seq data depicting STAT5 binding, H3K27ac patterns and RNA-seq data from the *Wap* gene in L1 mammary tissue. (B) Close-up of constituent enhancers E1 and E2. The region flanking both enhancers is enriched for Pol II binding. RNA-seq and RACE data demonstrate that E1 is transcribed into several polyadenylated eRNAs of varying length, depicted with purple arrows. (C) Close-up of constituent enhancers E3 and S4. The region surrounding E3 is enriched for Pol II binding, while a single Pol II peak is observed under S4. RNA-seq and RACE data demonstrate that both regions are transcribed into polyadenylated eRNAs, either uni-or bidirectionally. Enhancer RNAs are depicted with violet and green arrows (E3 and S4 eRNAs, respectively). (D) Schematic depiction of enhancer mutants and *Wap* expression in L1 mammary tissue as determined by qPCR, relative to wild-type mice.

To confirm the presence of eRNAs and determine their exact sequence, rapid amplification of cDNA ends (RACE) was performed followed by Sanger sequencing. This revealed bi-and unidirectional transcription of polyadenylated eRNAs originating from enhancers E1, E3 and S4, ranging from 2.7 to 0.5 kb in length (Fig. 1B, C, S1). RNA-seq data showed that the antisense S4 eRNA (S4 eRNA1) was the most abundant eRNA within the super-enhancer, followed by E1 eRNAs (Fig. 1B, C). The eRNAs transcribed from the sense strand at E3 and S4 (E3 eRNA and S4 eRNA2) are equivalently abundant. Interestingly, eRNA levels and Pol II occupancy at this locus were not correlated with the degree of STAT5 occupancy. However, Pol II occupancy is correlated with the expression of eRNA at any given enhancer.

To determine the functional significance of each constituent enhancer, mice harboring deletions of the γ-interferon activating sequence of enhancers E1 (ΔE1), E2 (ΔE2) or E3 (ΔE3) as well as a combinatorial deletion of all three motifs (ΔE1/2/3) had been generated. The contribution of each enhancer to *Wap* expression was characterized (14) (Fig. 1D). Mice lacking S4 were generated by CRISPR/Cas9-mediated deletion, and no reduction in *Wap* expression was observed (15). These data demonstrate that the extent to which a constituent super-enhancer element is transcribed; i.e. the abundance of specific eRNAs, is not correlated with its capacity to activate expression of the target gene.

### *Coordinated eRNA expression within the* Wap *super-enhancer*

Next we investigated potential interactions between enhancers that might regulate their transcription. ChIP-seq data supported a critical role for E3 in the recruitment of STAT5 to all enhancers within this locus, as ΔE3 mice showed reduced STAT5 binding at all other enhancers (14) (Fig. 2A). Given the dominance of E3 within the super-enhancer, we investigated its ability to regulate the expression of eRNAs at each constituent enhancer. Taqman probes were designed to measure the expression of E1, E3 and S4 eRNAs. Mice lacking all three enhancers showed an average reduction in transcription of 95% at each constituent enhancer (Fig. 2B). In ΔE3 mice, the average reduction was approximately 75%, while the transcription of each enhancer in ΔE1 mice was 50% that of wild-type on average, although this was not statistically significant. Loss of E2 only resulted in a significant decrease of E1 eRNA, resulting in approximately 50% of wild-type expression. Thus, the reduction in eRNA and *Wap* expression in these mice show an identical pattern (Fig. 1D, 2B). This demonstrates that eRNAs levels associated with constituent elements of a super-enhancer are indicators of overall super-enhancer activity, but not the activity of each individual enhancer.

**Figure 2.**
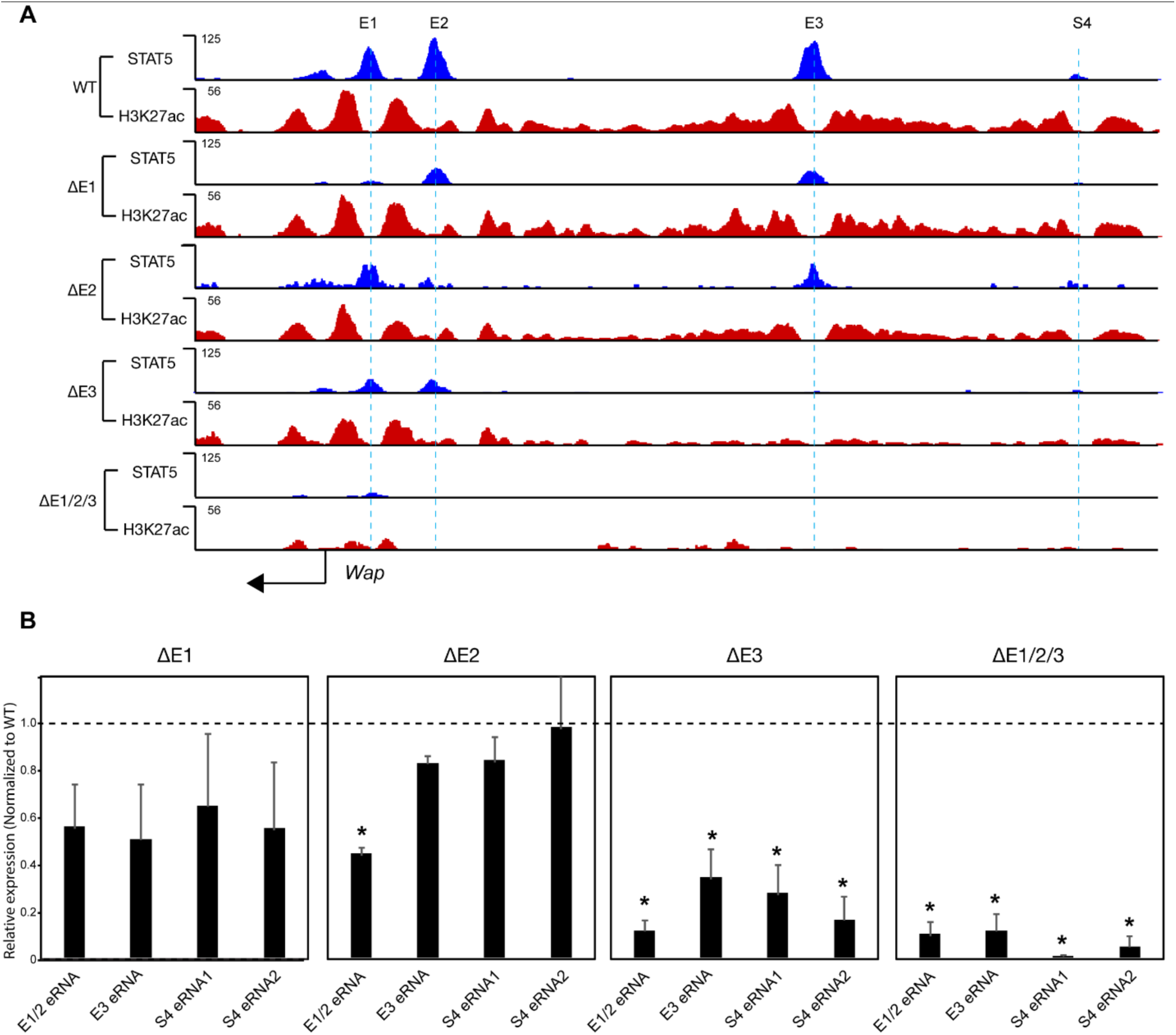
Deletion of STAT5 binding sites in constituent elements of the *Wap* super-enhancer result in a reduction of STAT5 binding, H3K27Ac, and enhancer transcription. (A) ChIP-seq from L1 mammary tissue in mice bearing individual or combined enhancer deletions demonstrates cooperativity among enhancers to effectively recruit STAT5. Loss of all three enhancers abolishes H3K27Ac at the locus. (B) Expression of eRNAs in wild-type and mutant mice L1 mammary tissue was measured by Taqman assays, and Ct values were normalized to *Gapdh* (wild-type, n = 10; ΔE1, n = 3; ΔE2, n = 2; ΔE3, n = 4; ΔE1/2/3, n = 6). Combined loss of E1-3 reduced expression of all eRNAs by approximately 90%, while loss of E3 alone reduced eRNA expression approximately 60-90%. Loss of E1 and E2 did not result in a statistically significant reduction of eRNAs originating from any enhancer, other than those deleted. Student’s t-test was used to evaluate changes in gene expression between mutant and wild-type mice. * p < 0.05

### *Constituent enhancers serve as transcription start site for* Ramp3 *in the absence of an endogenous promoter*

Although alternative promoter usage by both upstream and intronic enhancers has been widely reported (16-19) it is not clear whether they can serve as promoters for juxtaposed non-target genes. We therefore investigated the possibility that constituent elements within the *Wap* super-enhancer could serve as alternative promoters for the *Ramp3* gene, located 11 kb downstream of S4. To address this question, we generated mice carrying deletions between S4 and the first intron of *Ramp3* (MutA, MutC-E) and between E3 and the first intron (MutB) (Fig. 3, S2). By deleting the *Ramp3* promoter and upstream regions, we found that both E3 and S4 can serve as transcription start sites for *Ramp3* in the absence of the endogenous promoter. Furthermore, several splice forms were observed in mutants that use E3 and S4 to initiate *Ramp3* transcription (Fig. 3, S2, S3). This demonstrates that transcribed enhancers can be “coerced” into serving as transcription start sites for a coding gene. Furthermore, the ability to do so does not require the acquisition of H3K4Me3 marks characteristic of promoters (20). This obscures the distinction between enhancers and promoters, and raises questions concerning the nature of mRNA splicing.

**Figure 3.**
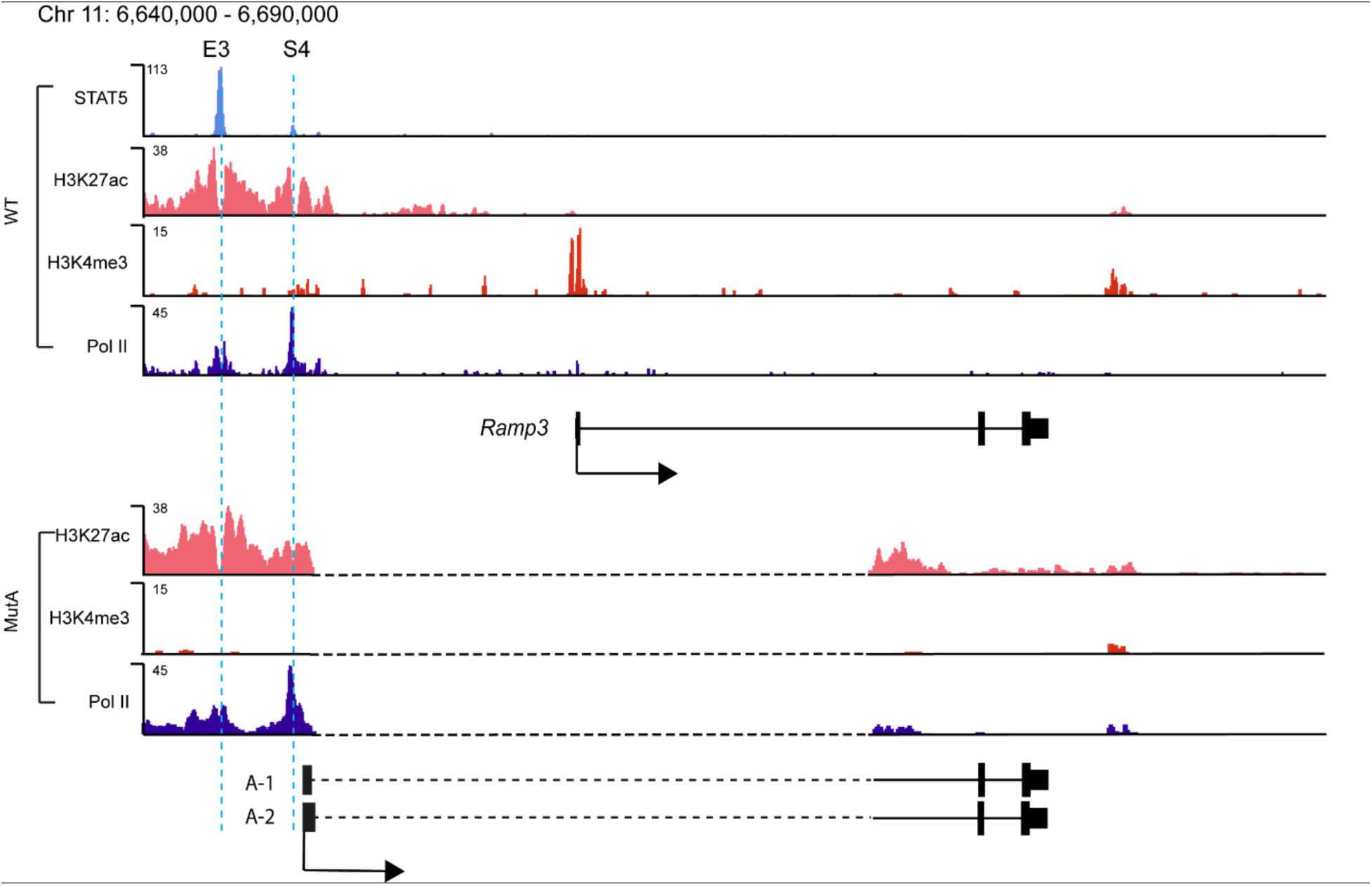
ChIP-seq in wild-type and MutA mammary tissue at the *Ramp3* locus. Mice harboring a deletion (depicted with dotted line) between the second *Ramp3* exon and the *Wap* super-enhancer utilized S4 as an alternative transcription start site, resulting in two splice forms. Utilization of S4 as an alternative transcription start site for Ramp3 did not require the acquisition of H3K4Me3 marks characteristic of promoters.

## Discussion

Genome-wide studies have provided observational data demonstrating that the establishment of activity-regulated enhancers, e.g. those established in response to LPS or estradiol treatment, is accompanied by their transcription into eRNAs (2,4,13,21). This has led to the development of algorithms that predict enhancers based on their production of eRNAs (12,13). In fact, it has been suggested that synthesis of eRNAs is more indicative of enhancer activity than H3K27Ac coverage (12). Thus, eRNAs have become widely accepted as a surrogate for enhancer activity, although it remains controversial whether eRNAs are a causal to enhancer-mediated gene activation, or merely a byproduct of Pol II binding (22). Knockdown experiments have implicated eRNAs in facilitating enhancer-promoter looping as well as promoting chromatin accessibility and Pol II recruitment (6,7). A recent study suggests a role for eRNAs in regulating the enzymatic activity of CBP/p300 at their associated enhancers (23). However, such functions have not been observed at an exceptionally active mammary super-enhancer, where ablation of the highly expressed S4 eRNA by genetic deletion neither reduced H3K27ac nor target gene expression. Together with previous studies, these results provide genetic evidence that eRNAs are not required for the induction of a target gene expression (10,11). It is more likely that in most cases transcripts originating from enhancers are mere byproducts of TF-mediated Pol II binding. Moreover, our data show that eRNAs are not necessarily accurate representations of enhancer activity. In fact, the putative enhancer element associated with the most abundant eRNAs and strongest Pol II binding had no visible impact on expression of the target gene when deleted. Thus, caution should be applied when using histone marks and transcription activity to predict enhancers within the genome.

Long non-coding RNAs (lncRNAs) and eRNAs are primarily distinguished by the ratio of H3K4Me1 to H3K4Me3 at their transcription start sites, as well as the observation that lncRNAs are more frequently spliced (3,24). It has been shown that H3K4Me3 marks facilitate recruitment of the U1 splicing machinery (25). Thus, the lack of this histone modification at enhancers could explain the lack of observed eRNA splicing (20). However, the STAT5 binding site S4 served as transcription start sites for *Ramp3* in the absence of its endogenous promoter, resulting in multiple splice forms without the acquisition of H3K4Me3 marks. It has previously been shown that intragenic enhancers can serve as alternative promoters, but the change in histone modifications at the alternative promoter were not characterized (19). It has also been suggested that eRNAs are rarely spliced due to the depletion of U1 splice sites near enhancers (26,27), yet several branch points exist near the *Wap* super-enhancer units E3 and E4 that are utilized when chimeric enhancer-*Ramp3* RNA is transcribed. We propose that hot spots of splicing machinery exist on the genome. When enhancers are placed close to these hot spots, the transcriptional machinery recruits splicing machinery and produces spliced eRNAs. Such spliced and polyadenlyated non-coding RNAs have been identified in mammary tissue carrying non-productive viral LTRs (28).

This study demonstrates that eRNA levels are not representative of enhancer activity but rather reflect the degree of Pol II occupancy. Our findings also raise questions regarding the regulation of splicing and demonstrate a need to reconsider the criteria that distinguishes eRNAs from lncRNAs. It could be argued that the distinction of eRNAs and lncRNAs is merely semantic and only depends on whether Pol II induced transcription encounters transcriptional termination signals before it is subject to splicing. Recent reports have shown that lncRNAs *Linc-p21* and *Lockd* are genuine eRNAs, and these findings suggest that distinguishing the two requires experiments that unlink a given transcript from the underlying DNA sequence (10,11,29).

## Methods

### Generation of mutant mice by CRIPSR-Cas9

CRISPR sgRNA constructs were designed based on their proximity to the mutation sites and their off-target scores (calculated by the online tool at crispr.mit.edu). Four sgRNAs 5’-TGAGTTTCCCATATGAACTC -3’; 5’-TCCTGATGGCCCTCTCCTGC-3’; 5’ – GCTCAGTGGAGAACCAATCT -3’ and 5’-TGGGTGTATGTAAGTAAGTG -3’ were cloned into the pDR274 vector (Addgene #42250) separately, and injectable RNAs were *in vitro* transcribed using the MEGAshortscript T7 kit (Life Technologies). Cas9 mRNA was *in vitro* transcribed from plasmid MLM3613 (Addgene #42251) using the mMESSAGE mMACHINE T7 kit (Life Technologies).

Mice used in this study were handled and housed in accordance with NIH guidelines. Animal experiments were approved by the NIDDK Animal Care and Use Committee. Zygotes preparation and microinjection were performed as previously described (30). Superovulated B6CBAF1/J female mice (JAX) were mated with B6CBAF1/J males, and fertilized eggs were collected from their oviducts. For microinjection, 100ng/µl of Cas9 mRNA and 50 ng/µl of each sgRNA in nuclease-free microinjection buffer (10 mM Tris, pH 7.5, 0.1 mM EDTA) were microinjected into the cytoplasm of fertilized eggs. Injected zygotes were cultured overnight in M16 medium at 37°C in 5%CO2. The next morning, those embryos that had reached the 2-cell stage were implanted into oviducts of pseudopregnant fosters (Swiss Webster, Taconic Farm). Genotyping was done to get the homozygous mice.

### RACE

Cleaned up RNA was used in a RACE assay using the FirstChoice RLM-RACE Kit (Ambion) according to the manufacturer’s instructions. RACE-specific primers had a final concentration of 0.2 ng/µL (reaction volume of 50 µL) and were synthesized by Eurofins Scientific. RACE-PCR components were replaced by ‘LongAmp Tag 2X Master Mix’ (New England BioLabs). Cycle conditions were: 35x (94°C 30”, 63°C 30”, 72°C 7’30”), 1x (72°C 7‘). The nested RACE-PCR was performed under unaltered conditions. The following primers were used:

<u>5’RACE</u> – *S1/2 Downstream,* Outer Primer 5’-GCAAGATGCAGAGAGAACAGAGC-3’; Inner Primer 5’-TACTGCCATGCTTGTGTTAGTC-3’.

*S3 Downstream,* Outer Primer 5’-CTCTTCCACCCTGTCCACTGCTC-3’; Inner Primer 5’-AGAGTTGATGGGGCAGGAAAGAGCC-3’.

*S4 Upstream*, Outer Primer 5’-TACACTTGGGCATCAGAGTCTGCC-3’; Inner Primer 5’-GTCAGCGCCTCTCCTCAGTAAACC-3’.

<u>3’RACE</u> – *S1/2 Downstream,* Outer Primer 5’-ACATTTAGGCCACTGTCAGG-3’; Inner Primer 5’-CCTTAGATGGGGATGACTGCC-3’.

*S3 Downstream*, Outer Primer 5’-CACATAGTAGCCGAGGATGGCC-3’; Inner Primer 5’-GGCACCTGCCTCCTCCTTCTAGTCT-3’.

*S4 Upstream*, Outer Primer 5’-GCTCAGCAGCACCAAGGGTAAT-3’; Inner Primer 5’-CCAAAGCTCCTTGGCCTGTC-3’.

*S4 Downstream*, Outer Primer 5’-GGAGGAGTTGTTTGGCTTCATGC-3’; Inner Primer 5’-GGAAGCTGTTAGTTACCGGTCAAGC-3’.

Samples were loaded to a 1% agarose gel (100V 50’) and visible bands were purified from the gel using MinElute Gel Extraction Kit (Qiagen). Sequencing results were provided by Macrogen.

### qRT-PCR

Total mammary tissues were harvested at lactation day 1 followed by RNA extraction. RNA was extracted using the PureLink RNA Mini Kit (Ambion) according to the manufacturer’s instructions. Complementary DNA (cDNA) was synthesized from total RNA using Superscript II (Invitrogen) with random hexamers and quantitative PCR was performed with the following Taqman probes.

*E1/2 eRNA*, Forward: 5’-CACCTCTCTGCTCTGTTTCTAC-3’; Reverse: 5’-GAACTGAAGCACTGGCAATTT-3’; Probe: 5’-ATGAGTTCCGGCAGCCTGTTGA-3’.

*E3 eRNA*, Forward: 5’-CAAAACCCTGGTGTCCTTTTC-3’; Reverse: 5’-TGTGCTTCCAGTATGCGTAG-3’; Probe: 5’-TTTCCTGCCCCATCAACTCTGCT-3’.

*S4 eRNA1*, Forward: 5’-AATCAAACCCGTTCCTCTGG-3’; Reverse: 5’-AGTCTCTTGCAGAAAGCCTG-3’; Probe: 5’-AGCCCCTTTCACCTTCTAACATCCAC-3’. *S4 eRNA2*, Forward: 5’-GCATGTTCTGAACATGGCTTGC-3’; Reverse: 5’-TGGCTTCTAGATGGTTCCCAAAGA-3’; Probe: 5’-AAGACAGCCAGTTTGGATCAGCCA-3’.

Real-time PCR was carried out using the BioRad CFX96 Real-Time PCR Detection System (185-5196; BioRad). Individual PCRs were performed in triplicate using the reference gene GAPDH for normalization. The threshold cycle (Cq) was determined by default settings. Relative gene expression (as a fold change) was calculated with the 2-*ΔΔ*Cq method. Data were presented as standard deviation in each group and were evaluated with a two-tailed, unpaired Student’s *t*-test using PRISM GraphPad. Statistical significance was obtained by comparing all groups to wild type. A value of P < 0.05 was considered statistically significant.

### ChIP-seq

Frozen-stored mammary tissues were ground into powder and then crosslinked with 1% formaldehyde (Sigma-Aldrich) for 10 min. Fixed nuclei were isolated, followed by chromatin fragmentation using sonicator 3000 (25 cycles; 30 sec pulse/30 sec rest, Misonix Sonicators). One milligram of chromatin was immunoprecipitated with Dynabeads Protein A (Novex) coated with anti-H3K4me3 (Millipore, 17-614), anti-H3K27ac (Abcam, ab4729), anti-RNA Polymerase II (Abcam, ab5408) and anti-STAT5A (Santa Cruz, sc-1081). After serial bead washes, ChIPed DNA was reverse-crosslinked and purified. The DNA fragments were blunt-ended using End-it DNA End-Repair Kit (Epicentre Biotechnology), ligated to the Illumina Indexed DNA adaptors, and sequenced with HiSeq 2000 (Illumina).

### Data analysis

ChIP-seq data were mapped to the reference genome mm10 using Bowtie aligner (31) and visualized using HOMER (http://homer.salk.edu/homer/) (32) and IGV (33). For visualization, the total reads number of mapped result in each sample was normalized to 10 million and background signals of less than 2 were eliminated. All the ChIP-seq data generated in this study were deposited in GEO database. ChIP-seq datasets of wild-type and *Wap* enhancer mutant mammary gland were obtained from GSE74826.

#### Acknowledgements

The authors thank Chithra Keembiyehetty Nightingale, Sijung Yun and Harold Smith from the NIDDK Genomics Core for NGS and Chengyu Liu from the NHLBI mouse core for generating mutant mice.

## Author Contributions

CW designed experiments, supervised the project, conducted RACE and ChIP-seq studies, analysed data and wrote the manuscript. TMK identified founder mice, established lines, conducted expression studies and wrote the manuscript. MW analysed ChIP-seq and RNA-seq data. CR conducted RACE and expression studies. LH conceived, designed and supervised the study, analysed data and wrote the manuscript.

